# DNA nullomers as a statistical phenomenon

**DOI:** 10.64898/2026.10.10.758214

**Authors:** Luc Wieners, Martin E. Garcia

## Abstract

Nullomers are sequences absent from a certain genome. Due to their unexpected occurrence in short sequences, a hypothesis is that they are avoided for evolutionary reasons or because they might have lethal consequences on organisms. However, we present evidence that the occurrence and frequency of nullomers in genomes is expected as a stochastic consequence of uneven base pair combination distributions in the genetic code. This is done by artificially constructing whole genomes from statistics with a Markov chain model. The number of nullomers in these artificial genomes is in good agreement with the nullomer counts in reference genomes. Additionally, we investigated the nullomers of over 200 human genomes and found large deviations in the nullomers between individuals, further supporting the hypothesis that a vast majority of nullomers is very unlikely to have lethal effects.

## Introduction

An important step to understanding genomes better is to investigate which sequences do not occur in them. Due to the non-existence of these sequences, they have been called “nullomers” [1]. This concept is linked to bioinformatics, where “minimal absent words” are defined as the shortest strings (or “words”) not included in a larger string (for example a genome) while all substrings are included. Therefore, the origin of nullomers and especially minimal absent words and their computation [2–7] originally lies in biocomputing.

However, the idea of non-existent sequences in genomes is also of great interest for biological applications, leading to their study in more applied contexts [8, 9]. These applications of nullomers [10] cover a wide range from detecting cancer [11] over developing peptides [12] with anti-cancer properties [13–15] to the use as tags in forensics [16]. Furthermore, nullomers are studied in viruses—namely, Sars-CoV2 [17], influenza [18] and Ebola [19] – and they are used in phylogeny [20–22] as a distance measure between species.

A remaining question is the origin of nullomers. Do they occur through natural selection, could they even be lethal or are there other reasons for sequences being absent from a genome? One important characteristic of nullomers has been identified: They contain an over proportional amount of CpG dinucleotides [23–26], which is usually quite rare in genomes due to their high mutation rates [27–30]. Furthermore, it has been investigated which nullomers are “significant”, i.e. they are expected to occur but are instead absent in a genome [31].

In this work, we continue the investigation of the reasons behind the occurrence of nullomers. To do so, we use Markov chain models [31, 32] to create artificial genomes [32]. These genomes are created for different species and the nullomers and their counts are computed for the generated genomes and compared with reference genomes. Additionally, we investigate the differences between the nullomers of a large set of individuals. The differences of minimal absent words in four human genomes have been investigated in 2011 [33]—now, due to the fast advances in DNA sequencing, many high coverage genomes are available. We therefore computed the nullomers of the 232 human genomes of the human pangenome project which aims to create a diverse representation regarding human genomes [34].

From these investigations, we find that artificial genomes have nullomer occurrence and counts which are in good agreement with reference genomes. Furthermore, there is a large degree of variation between the nullomers of different human genomes. These findings imply that the majority of nullomers occurs for statistical reasons which lie in the uneven distribution of the genetic code rather than by natural selection of certain sequences.

## Results

### Nullomers among different species

Since genomes show a high degree of variation regarding their length and composition, we investigated the genomes of ten different species. These species will then be used for the comparisons between reference and artificial genomes.

For these species, we selected the lungfish (*Lepidosiren paradoxa*) genome, as it is the longest known animal genome, at nearly 90 gigabasepairs (Gbp)—around 30 times the length of the human genome. Then, we chose the human (*Homo sapiens*) genome, since human genomes are studied in detail in the later sections, and the mouse (*Mus musculus*) genome as a model organism with a genome length similar to humans. Additionally, we selected fruit fly (*Drosophila melanogaster*) and thale cress (*Arabidopsis thaliana*) as two model organisms with medium-sized genomes and to also include a plant. Furthermore, we chose plasmodium (*Plasmodium falciparum*), yeast (*Saccharomyces cerevisiae*), streptomyces (*Streptomyces coelicolor*) and E. coli (*Escherichia coli*) as shorter genomes with differing GC content (the combined amount of guanine and cytosine in a genome), namely 19 %, 38 %, 51% and 72 %, while also including yeast and E. coli as model organisms. Finally, we selected the corona virus Sars-CoV2 as an example for a very short genome. The sequences are taken from the NCBI [37] assemblies page with the accession codes listed in the Methods section “Genome selection”.

For this diverse set of species, the amount of nullomers is computed for sequence lengths between one and 15. As it is commonly the case, a nullomer is here defined as a DNA sequence which is absent from the whole genome, including the reverse complement strand. The procedure of the computation of nullomers is described in the Methods sections “Genome reading” and “Nullomer and sequence counting” where the computer code is reviewed in detail. In Table 1, the amount of nullomers for sequence lengths up to 15 is written for the investigated species. We can see that the shortest sequence length at which nullomers occur is six (for streptomyces and corona), whereas the latest occurrence is observed at length 12 for the lung fish genome.

**Table 1.** Nullomer counts among species. For the ten selected species in this study, the amount of nullomers for sequence lengths up to 15 is given. Species are ordered by genome length.

| Species name<br>(common name) | Length 6 | Length 7 | Length 8 | Length 9 | Length 10 | Length 11 |
| --- | --- | --- | --- | --- | --- | --- |
| *Lepidosiren paradoxa*<br>(lung fish) | - | - | - | - | - | - |
| *Homo sapiens*<br>(human) | - | - | - | - | - | 104 |
| *Mus musculus*<br>(mouse) | - | - | - | - | - | 188 |
| *Drosophila melanogaster*<br>(fruit fly) | - | - | - | - | - | 116 |
| *Arabidopsis thaliana*<br>(thale cress) | - | - | - | - | 86 | 22,858 |
| *Plasmodium fal-*<br>*ciparum*<br>(plasmodium) | - | - | 146 | 9,354 | 178,435 | 1,737,522 |
| *Saccharomyces*<br>*cerevisiae*<br>(yeast) | - | - | - | 2 | 6,455 | 463,242 |
| *Streptomyces coe-*<br>*licolor*<br>(streptomyces) | 1 | 26 | 1,296 | 30,122 | 329,145 | 2,318,456 |
| *Escherichia coli*<br>(E. coli) | - | - | 52 | 2,502 | 68,553 | 1,269,966 |
| Sars-CoV2<br>(corona) | 127 | 3,218 | 34,489 | 214,336 | 992,746 | 41,356,66 |
| **Species name**<br>**(common name)** | **Length 12** | **Length 13** | **Length 14** | **Length 15** |  |  |
| *Lepidosiren para-*<br>*doxa*<br>(lung fish) | 183 | 76,134 | 3,990,901 | 86,552,338 |  |  |
| *Homo sapiens*<br>(human) | 44,233 | 2,346,390 | 40,780,234 | 405,269,832 |  |  |
| *Mus musculus*<br>(mouse) | 53,122 | 2,568,458 | 42,584,571 | 413,950,050 |  |  |
| *Drosophila mela-*<br>*nogaster*<br>(fruit fly) | 179,270 | 11,452,426 | 147,059,268 | 895,139,174 |  |  |
| *Arabidopsis thali-*<br>*ana*<br>(thale cress) | 1,157,788 | 20,186,540 | 168,996,159 | 922,018,254 |  |  |
| *Plasmodium fal-*<br>*ciparum*<br>(plasmodium) | 11,253,242 | 56,982,298 | 252,716,616 | 1,052,268,984 |  |  |
| *Saccharomyces*<br>*cerevisiae*<br>(yeast) | 7,229,055 | 51,021,594 | 248,244,629 | 1,051,789,596 |  |  |
| *Streptomyces coe-*<br>*licolor*<br>(streptomyces) | 12,698,109 | 59,693,528 | 257,434,613 | 1,059,859,014 |  |  |
| *Escherichia coli*<br>(E. coli) | 11,081,557 | 59,403,364 | 259,828,234 | 1,064,817,366 |  |  |
| Sars-CoV2<br>(corona) | 16,717,797 | 67,049,242 | 268,375,768 | 1,073,682,114 |  |  |

Additionally, it can be seen that for even sequence lengths, the amount of nullomers is always an even number. This can be explained by the fact that, if a sequence is a nullomer, its reverse complement is also a nullomer, since both DNA strands are considered during the search. Consequently, each nullomer is paired with a second nullomer. While this holds for all sequence lengths, sequences of an even length can be palindromic, meaning that their reverse complement is the same as the original sequence (for example AACGTT). These sequences do not have a second, matching sequence which leads to odd nullomer counts. Sequences of an odd length, however, cannot be palindromic, since the base pair in the middle of the sequence would have to be matched up with its complement which leads to even nullomer counts.

For a better visualization, the logarithm of the nullomer counts is shown in Figure 1 for all species. We observe an inverse correlation between genome sizes and the sequence length at which nullomers start to occur. This is expected, since a longer genome means that there are more chances for a certain sequence to occur.

**Figure 1.**
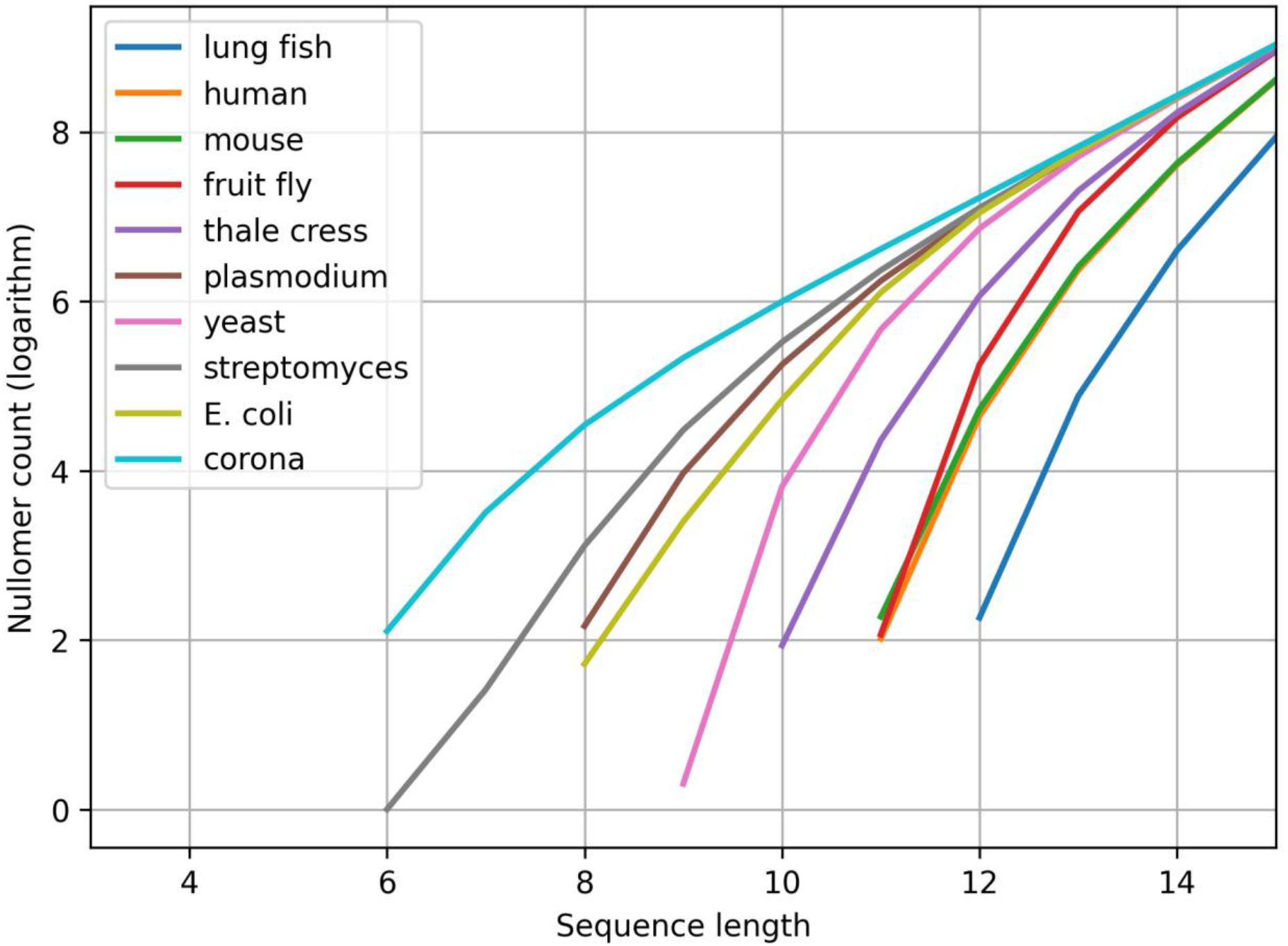
Nullomer distribution differences among species. For ten species, the logarithm (base 10) of the computed amount of nullomers is shown for sequence lengths up to 15. The graphs start at the smallest sequence length for which nullomers occur for the respective species. The order in the legend is based on the genome length.

Furthermore, it can be observed that there is a limit which is approached by all graphs which corresponds to almost all sequences being nullomers. This is the expected limit case for longer sequence lengths since the number of possible sequences grows exponentially. The described limit case is a linear function in the logarithmic plot in Figure 1. It can be seen that this line is approached first by the species with smaller genomes, as expected. For example, for sequence length 15, the number of possible combinations is 1,073,741,824 which is very similar to the amount of nullomers for the five species with smaller genomes.

### Artificially constructed genomes

For the main goal of this work—assessing, if the nullomer counts in nature are expected from statistics alone—we consider “artificial genomes”. These genomes are created via a Markov chain, a probabilistic model which predicts a future state based on a previous state and transition probabilities. The Markov chain model can be applied to any sequence and is therefore also used for DNA sequences [31] and generating genomes [32].

It works by considering a sequence A with *n* nucleotides and predicting a follow-up sequence B with *m* further nucleotides (here, we describe such a Markov chain as *M*(*n, m*)). This prediction is based on transition probabilities: B is predicted as a follow-up of A with a probability *P*_A_(B). Consequently, these in total *n* · *m* probabilities have to be known for the model to work. In most cases, they can be computed from a given data set – in this case, they are determined from the given genome of the respective species. With these transition probabilities computed, it is then possible to generate a whole genome with the same length as the reference genome. The exact procedures used for these calculations are described in the Methods section “Genome generation”.

In this work, we generate ten artificial genomes for each species to account for the stochastic nature of the method. The used Markov chains for the species are *M*(3,3) for the lung fish, human and mouse genome; *M*(5,1) for the fruit fly and thale cress genomes, *M*(3,1) for the plasmodium, yeast, streptomyces and E. coli genomes and *M*(2,1) for the Sars-CoV2 genome. It is important to choose a Markov chain with *n* and *m* as small as possible, since otherwise the information about nullomers could be stored within the computed transition probabilities. Therefore, the number of transition probabilities should be significantly smaller than the number of sequences of the length where nullomers start to occur. For example, for the Markov chains *M*(3,3) used for the human genome, there are 4,096 probabilities and nullomers start to occur at sequences lengths 11, which corresponds to 4,194,304 total sequences at this length. For this reason, the Markov chain type is varied based on the genome length. A prediction length of 1 is used for most genomes, the input length is varied based on the genome size. For the longer genomes with lengths over 1 Gbp, a prediction length of 3 was used with an input length of 3. In the cases of the thale cress and fruit fly genomes, *M*(3,3) leads to an underestimation of nullomers whereas for the mouse and human genome *M*(5,1) leads to an overestimation of nullomers. Differences in the genomes that could explain this behaviour are discussed in the section “Sequence spectra comparisons”.

Next, the number of nullomers is calculated for the ten artificial genomes per species in the same way as for the reference genomes. The resulting nullomer counts for the real and generated genomes are visualized in Figure 2. Over all ten species, a good agreement between the nullomer counts of real and artificial genomes is observed. Both overestimations of the number of nullomers (for example for the mouse genome) as well as underestimations (for example for the lung fish genome) are found.

**Figure 2.**
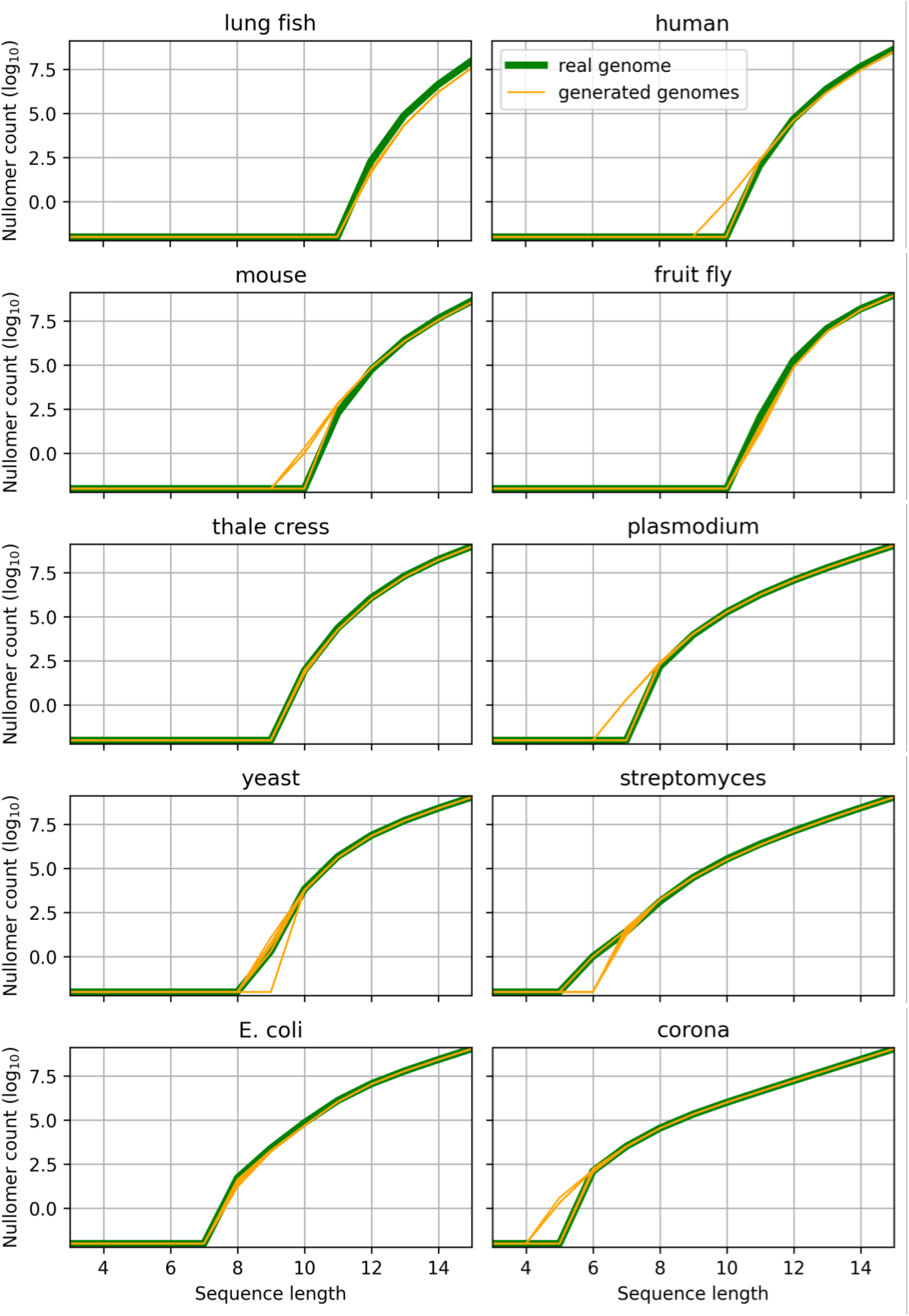
Nullomer counts in real and generated genomes. For the ten investigated species, the logarithm (base 10) of the number of nullomers at sequence lengths up to 15 is shown for both the real (green) and for ten generated genomes (orange). Nullomer counts of zero are displayed as –2 to avoid negative infinity due to the logarithm.

Additionally, it can be compared at which sequence lengths nullomers start to occur. For four species, the predicted minimal nullomer length is in agreement with the real minimal nullomer length for all ten artificial genomes. For the other cases, an average was computed which is displayed in Table 2. The rounded average is in agreement with the minimal nullomer length for eight species. Two exceptions can be found for the mouse genome, where six out of the ten artificial genomes predict the occurrence of nullomers at a sequence length of 10 instead of eleven, and for streptomyces, where the reference genome contains a single nullomer of length 6 but nine out of the ten generated genomes predict the minimal nullomer length as 7.

**Table 2.**
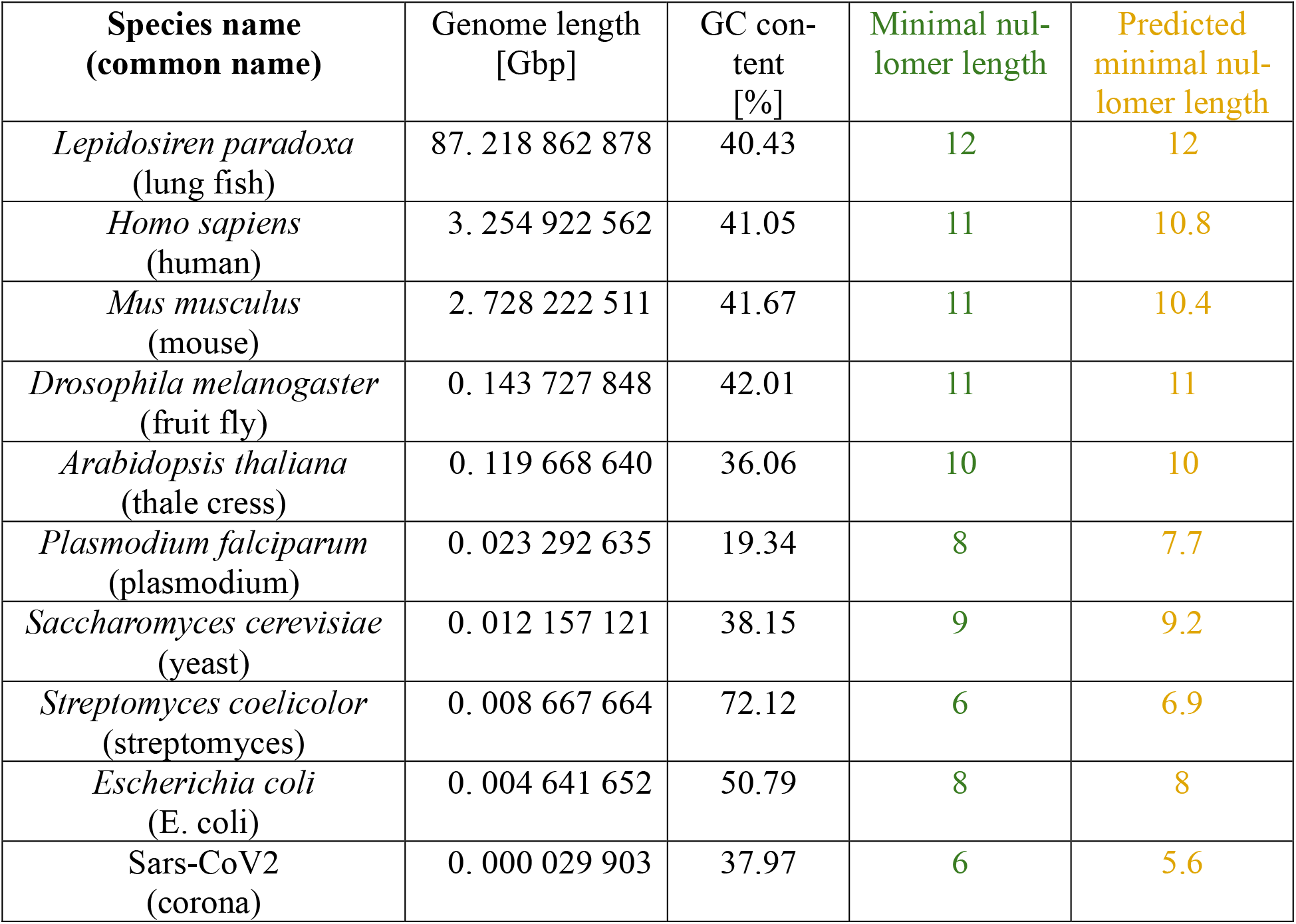
Minimal nullomer lengths among species. The length of the genome is given as well as the GC content for all ten investigated species. The minimal sequence length for which there are nullomers in a certain genome, is given for the reference genome and as an average for the predicted genomes using Markov chains.

In addition to the minimal nullomer lengths of reference and generated genomes, Table 2 also contains the genome length and the GC content for all of the investigated species. There, it can be seen that the minimal nullomer length decreases with the genome length, as discussed above.

However, there are two exceptions to this rule: plasmodium and streptomyces. While the plasmodium genome is longer than for example the yeast genome, it has a minimal nullomer length of 8, smaller than the yeast genome with a minimal nullomer length of 9. Similarly, streptomyces has a minimal nullomer length of 6—the same as the corona virus genome which is nearly 300 times shorter. Therefore, for both plasmodium and streptomyces, the occurrence of nullomers would be expected at longer sequence lengths.

An explanation can be found in the GC content of these genomes. For plasmodium, the GC content is exceptionally low at 19.34 % whereas it is extremely high for streptomyces at 72.12 %. This leads to a more uneven distribution of the nucleotide bases in these two genomes. As a consequence, nullomer occurrence is observed at lower sequence lengths, since there are more sequences which are more unlikely to be found in these two genomes. For plasmodium, sequences rich in C and G are therefore expected to occur much less and followingly become nullomers at shorter sequence lengths than in genomes with a more even distribution of the nucleotide bases. The same argumentation can be made in streptomyces for sequences that are rich in A and T.

### Sequence spectra comparisons

In addition to the investigation of nullomers, it is also possible to consider the counts of all sequences of a given length for the generated genomes in comparison with the real genomes. In Figure 3, this is visualized in the form of a map where the number of occurrences of a given sequence in the reference genome is plotted against the number of occurrences in the first generated genome. Due to large differences in the number of occurrences, both axes are logarithmical in base 10. Since there are over 4 million sequences of length 11, the visualization does not use individual points and is instead done via a colour map (also logarithmically in base 10) which represents the density of points.

**Figure 3.**
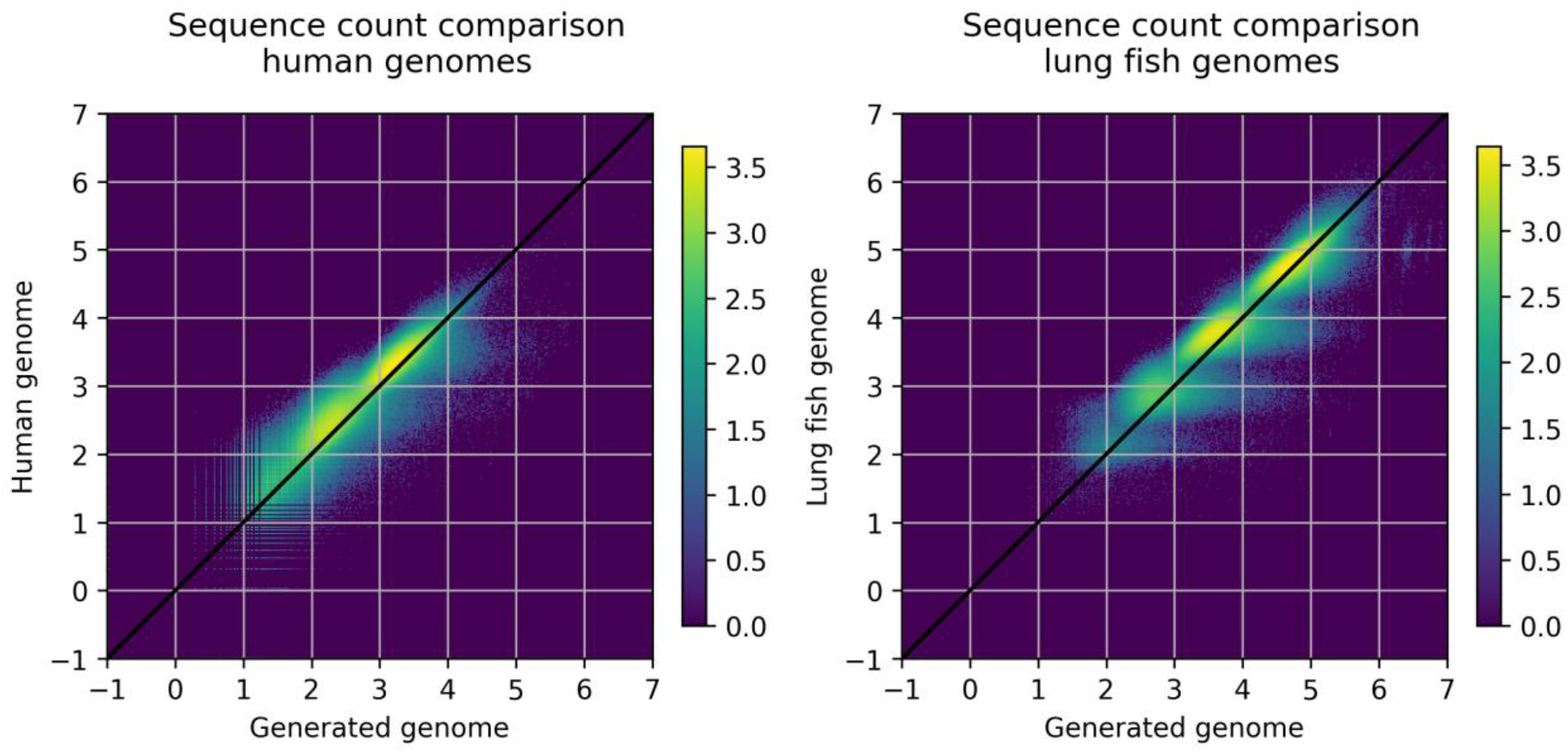
Sequence counts of length 11 in real and artificial genomes. The sequence counts for each sequence of length 11 of the reference genome are plotted against the sequence counts in a generated genome using a Markov chain—this is done for the human genome (left) and the lung fish genome (right). The plot is double-logarithmical in base 10 and the number of sequences in a region of the plot is indicated via colour (also logarithmical, base 10).

A reasonable agreement between the sequence counts in the real and the generated genome can be seen, meaning that the Markov chain approach can reproduce sequence counts of sequences longer than its input and output length. Upon closer inspection of Figure 3, it can be observed that the points indicating individual sequences are grouped into clusters. For the human genome, there are three of these clusters while the lungfish genomes have four.

Consequently, the number of sequences that appear *n* times does not follow a normal distribution in the human or lung fish genome but has maxima [35]. This behaviour has been called “multimodal” and is observed in the genomes of some species, mostly in tetrapods [32].

A visualization of the number of sequences which occur *n* times is shown in Figure 4 for the human and the lung fish reference genomes. There, the multimodal structure of the frequency spectrum can be seen clearly. Furthermore, the different maxima in this representation are known to be linked to the occurrence of the motif CG [32]. Since this motif is very abundant in nullomers, these frequency spectra also illustrate the occurrence of nullomers based on the usage of this motif.

**Figure 4.**
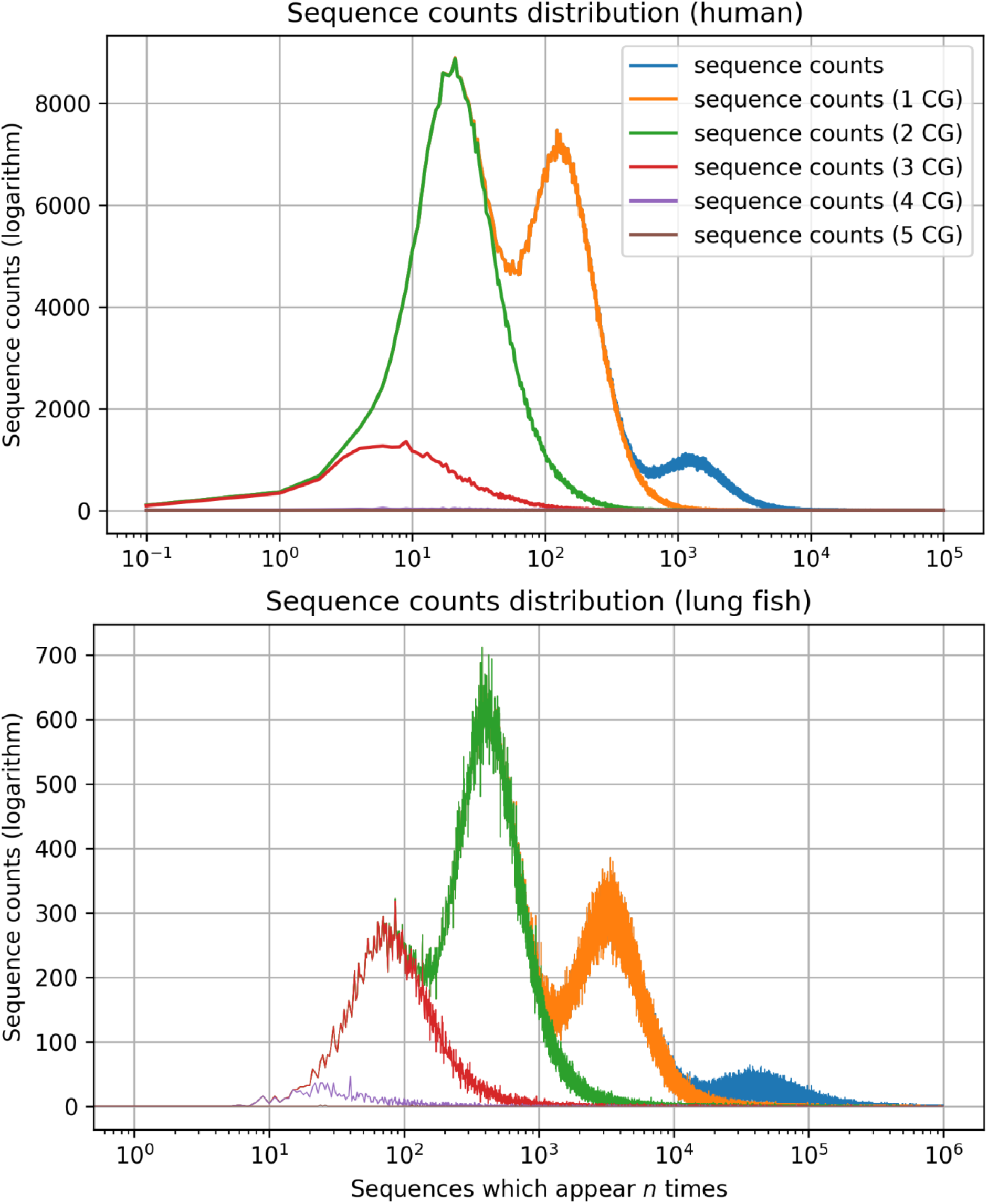
Frequency spectrum of 11-mers in the human and lung fish genome. For the human genome (top) and the lung fish genome (bottom), the number of sequences which occur *n* times are plotted against *n*. This can also be seen as the projection of Figure 3 onto the y-axis. Nullomers in the human genome are plotted at a x-value of 0.1; the lung fish genome has no nullomers of length 11. The curves overlap when they transition into each other (blue follows orange, orange follows green and so on).

It has been shown that the different maxima in multimodal frequency spectra are related to the different occurrences of the CpG dinucleotide [32]. Sequences containing one CpG motif are much less common than sequences with no CpG; sequences which contain the motif CG twice are even less common and so on. This behaviour is explained to the CpG suppression in these genomes which takes place due to the high mutation rates [27–30].

In Figure 4, the frequency spectrum of length-11-sequences with zero to five occurrences of the CG motif are shown separately. If sequences become less common, the amount of the short sequence CG increases.

Most of the human nullomers contain three copies of the CG motif. For the lung fish genome with nearly 90 Gbp, the spectrum is shifted to the right in comparison with the human genome since sequences on average occur more often. This leads to the formation of a fourth maximum, which is not present for the human genome, with sequences that contain the CG motif four times. No nullomers occur at length 11 since the spectrum is shifted towards higher occurrence numbers.

Additionally, Figure 4 demonstrates how the suppression of the CG motif leads to the formation of nullomers. If the CG motif was not suppressed, the sequence distribution would be a normal distribution at the position of the blue part of the curves. Consequently, sequences that appear less often would be rarer since the other maxima would be missing. Due to the suppression of the CpG dinucleotide, certain sequences get avoided which leads to some sequences occurring much less or even not at all.

This agrees with the observation that the nullomer occurrence length in *Drosophila melanogaster* (fruit fly) is at 11, the same as for the human genome even though the latter is over twenty times longer. Therefore, in comparison one would expect nullomer occurrence to begin at higher sequence lengths in the human genome. However, contrary to the fly genome, the human genome has strong CpG suppression which leads to a more uneven sequence distribution and therefore a lower nullomer occurrence length.

### Nullomer differences among human genomes

As a final test of the statistical factors underlying the occurrence of nullomers, we investigated a large number of human genomes and their shared nullomers of length 11. The human genomes were taken from the human pangenome project [34] for a diverse representation of human genomes.

First, we study how the number of shared nullomers changes if more and more human genomes are considered. To begin with, we can consider one genome. In this case, the average number of nullomers of length 11 over all considered genomes is 108. If a random second genome is added, the average number of nullomers of length 11 present in both genomes is only 87. This procedure can be continued for more and more genomes. The change in the number of genomes is visualized in Figure 5. To account for the stochastic elements in this procedure, it is repeated 100 times.

**Figure 5.**
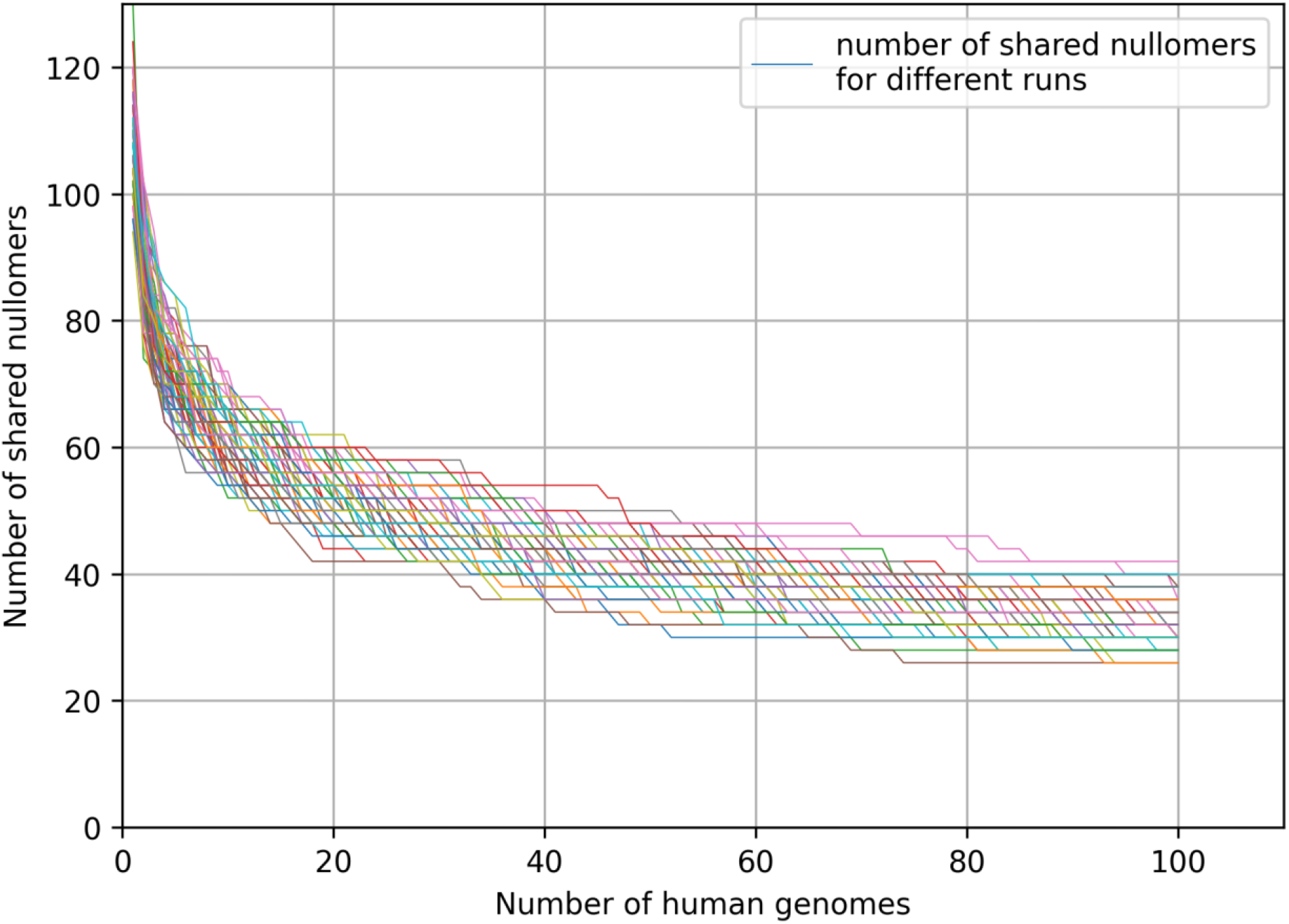
Shared nullomers among human genomes. The number of shared nullomers of length 11 between human genomes is shown for up to 100 genomes in 100 different runs (coloured lines). In each run, genomes are subsequently added to a list and the shared nullomers are computed at each step. The genomes are taken from the 232 genomes of the human pangenome project [34] and each genome is only used once per run.

By comparing the shared nullomers, we can observe that after including 100 genomes, the number of shared nullomers lies between 26 and 42, depending on the run. Note that there are only even numbers of nullomers due to the sequence length being odd—as discussed above. In addition to the low number of shared nullomers between 100 individuals, a continuous decrease is observed when more and more genomes are added. In summary, this demonstrates that there is a large degree of variation between the nullomers of length 11 in humans. Furthermore, variations are also found in the total number of nullomers of length 11 which ranges from 92 to 130.

If all 232 genomes from the human pangenome project are considered, the number of shared nullomers of length 11 is computed as 22 which equates to 20.4 % of the average number of nullomers for a single genome. Among all individuals, 304 sequences are a nullomer in at least one genome, almost three times as many as the average per genome.

In Table 3, all of the 22 shared nullomers (11 unique sequences if they are paired with their reverse complements) are listed. In addition to this, it was computed how often these sequences occur in the ten Markov-generated human genomes of the type *M*(3,3) and in ten other artificial genomes of the type *M*(6,3). The latter group with the longer input length into the Markov chain investigates the trend of the sequence counts if more information about the genome is available.

**Table 3.**
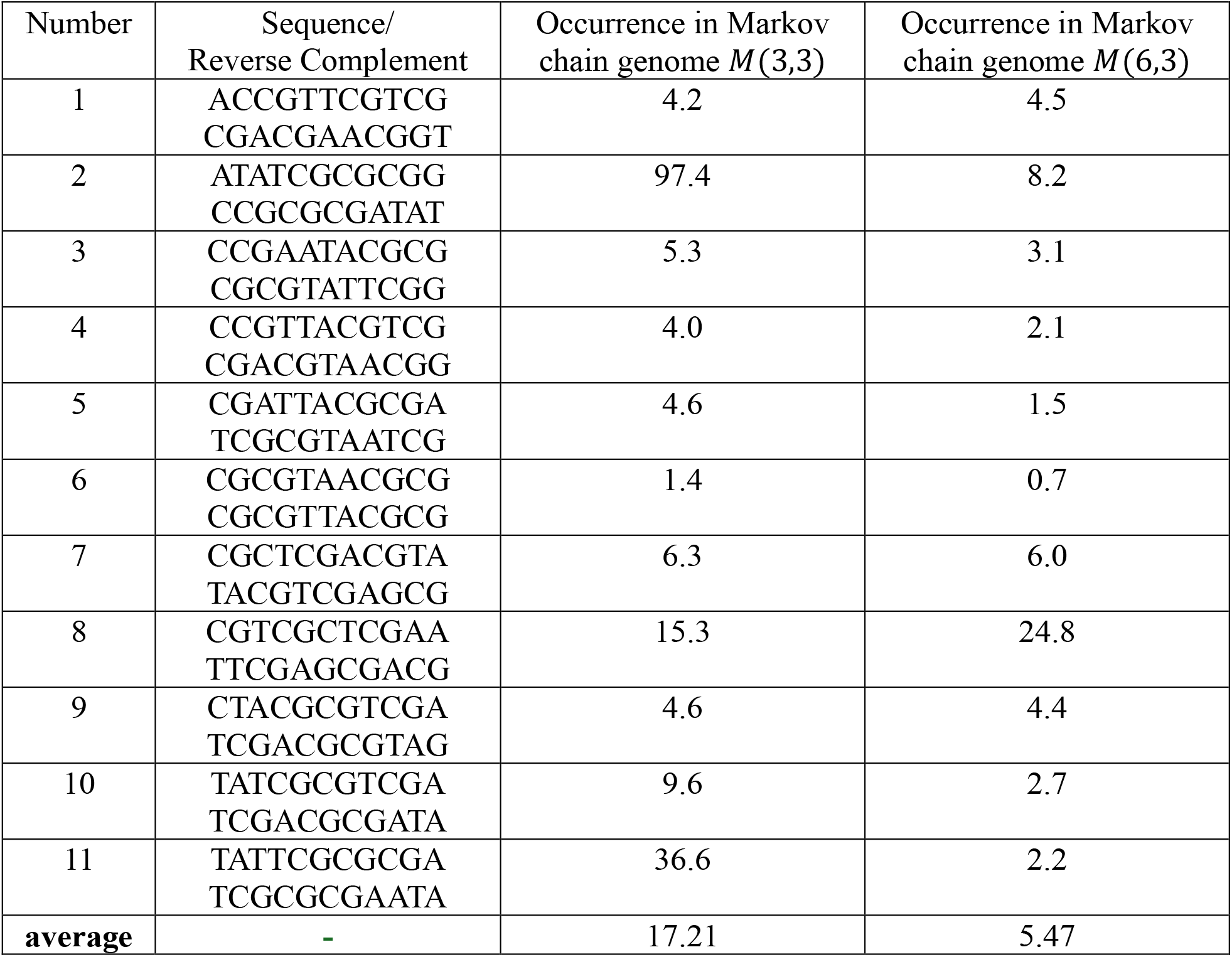
Shared nullomers among human genomes. The eleven nullomers of length 11 which were shared among all 232 human genomes from the human pangenome project are listed. Next to the sequence and its reverse complement, the average occurrences of these sequences in the artificial human genomes (generated via two different types of Markov chains) are shown for each sequence and on average.

Table 3 demonstrates that the sequences which are shared nullomers are also rare in the Markov genomes. Their average occurrence in the artificial genomes is below ten for eight sequences in the genome of the type *M*(3,3) and for ten sequences for *M*(6,3). The total average numbers of occurrences over all sequences (17.21 and 5.47) correspond to 1.09 % and 0.35 % of the average occurrences of a random sequence of length 11. A decrease in the occurrence counts from the genome *M*(3,3) to *M*(6,3) is found in nine sequences.

The significance of nullomers was investigated by Koulouras et al. [31] who computed which sequences are absent but highly expected to occur. A list of these sequences is provided on the Nullomers Database (nullomers.org) – for the human genome, 13 sequences are listed of which seven are unique and six are reverse complements (one sequence is a palindrome).

Here, these sequences are investigated within the 232 human genomes. In the following, it is listed in how many human genomes these 13 sequences occur.

– TATTATGCGCG/CGCGCATAATA: 14

– TTTCGCGAAATT/AATTTCGCGAAA: 5

– AAATTGGCGCAGG/CCTGCGCCAATTT: 0

– GGCGATTTTTGGG/CCCAAAAATCGCC 36

– TTTGGGCGCAACA/TGTTGCGCCCAAA: 5

– ATTTTTTACGGGC/GCCCGTAAAAAAT: 53

– ATTTCGCGAAAT: 0

To summarize, five of the seven sequences are present in some of the human genomes whereas two sequences are absent from all human genomes. Consequently, we observe that not even the nullomers that were evaluated as significant are all strictly absent among the 232 investigated human genomes.

## Discussion

We presented calculations that give evidence that the occurrence of nullomers can be explained with statistical models. Additionally, variation between nullomers among individual human genomes is observed.

These results support other work in which the occurrence of nullomers has been connected to statistics rather than natural selection or even toxicity [23, 31]. Nonetheless, there are still examples for sequences that are avoided due to natural selection, for instance the codon AGA in bacteria [38] which was linked to ribosomal stalling [39]. Furthermore, certain nullomer peptides with poly arginine tails were reported to have lethal effects on cells [14]. However, this is not in contradiction with the statistical explanation in this work since sequences whose absence can be explained with statistics can still have negative effects on organisms.

However, the occurrence of nullomers cannot be viewed as unexpected if the distribution of short sequences such as pairs of two codons are considered. Instead, the occurrence length and number of nullomers in humans can be approximately reproduced by using only the 4,096 transition probabilities between codons. An explanation for nullomers is thereby already found when considering much shorter sequences.

In addition to this, for nullomers of minimal length in human genomes, there exist large discrepancies between individuals. Not even the nullomers that are considered highly significant are all absent from the genomes of different individuals.

Therefore, an interpretation of absent short sequences as toxic or lethal to organisms is unlikely to be correct for the vast majority of nullomers. The agreement between nullomer counts in artificial and real genomes suggests that nullomers cannot be considered an unexpected phenomenon but are rather much expected to occur based on the distributions of very short DNA sequences such as codons. Similarly, very high or very low GC content in genomes and the suppression of the motif CG is linked to the occurrence of nullomers at low sequence lengths which supports the explanation that uneven distributions in genomes cause nullomers.

Future investigations with respect to the effects of different Markov chain parametrizations might be interesting. Also, time series forecasting with machine learning could be an alternative approach to study genomes. Additionally, it could be interesting to study if nullomer occurrence can be linked more strictly to codon usage biases.

To conclude, we demonstrated that the number and occurrence length of sequences absent from genomes (nullomers) can be predicted with statistical models. This indicates that the majority of nullomers can be explained from the uneven distribution in genetic codes such as usage biases of codons. In an extensive analysis of 232 human genomes, it was found that only a small fraction of nullomers of length 11 is shared among all individuals. Not even all nullomers that previously have been evaluated as significant are absent from the complete set of individuals. Due to good agreements in nullomer counts between real and artificial genomes, it has to be considered that nullomer occurrence cannot be viewed as a special phenomenon. For the example of the human genome, it is very expected that sequences of length 11 would contain nullomers if codon usage and transitions between codons are considered. Additionally, the missing consistency between genomes of a species demonstrates that a majority of nullomers cannot have lethal effects.

## Methods

### Genome selection

The genomes of the ten investigated species were taken from the Assemblies Database of the National Center for Biotechnology Information [37]. For nine species, the reference genome was used, with the exception of *Streptomyces coelicolor*, where no reference genome was available. In the following, a list with the accession codes of the used genomes is provided.

– *Lepidosiren paradoxa* (lung fish): Assembly: ASM4058144v1, GenBank: GCA_040581445.1

– *Homo sapiens* (human): Assembly: GRCh38.p14, GenBank: GCA_000001405.29

– *Mus musculus* (mouse): Assembly: GRCm39, GenBank: GCA_000001635.9

– *Drosophila melanogaster* (fruit fly): Assembly: Release 6 plus ISO1 MT, GenBank: GCA_000001215.4

– *Arabidopsis thaliana* (thale cress): Assembly: TAIR10.1, GenBank: GCA_000001735.2

– *Plasmodium falciparum* (plasmodium): Assembly: GCA_000002765, GenBank: GCA_000002765.3

– *Saccharomyces cerevisiae* (yeast): Assembly: R64, GenBank: GCA_000146045.2

– *Streptomyces coelicolor* (streptomyces): Assembly: ASM893130v1, GenBank: GCA_008931305.1

– *Escherichia coli* (E. coli): Assembly: ASM584v2, GenBank: GCA_000005845.2

– *Sars-CoV2* (corona): Assembly: ASM985889v3, GenBank: GCA_009858895.3

The 232 human genomes from the human pangenome project [34] were accessed via the Human Pangenome Data Explorer (https://data.humanpangenome.org/assemblies) using the Assemblies section. The second release was used and it was alternated between maternal and paternal haplotypes. Further processing of the genome files in the FASTA format is described in the following section.

### Genome reading

To use the various genomes for calculations, they have to be read in as FASTA files and stored in a numerical format. This was done straightforward by a function which reads through all characters in a text file line by line. In an array, the current base pair is saved as a number (0 for A, 1 for C, 2 for G, 3 for T). It has to be checked for both upper- and lower-case letters since both can occur in FASTA files. If no match is found, the array entry is written as –1; this can happen occasionally in some FASTA files where undetermined base pairs are designated with the letter N.

It has to be considered that the characters per line can vary from file to file. Additionally, a rough estimate of the genome size is necessary. The array is saved in the datatype “int8” to minimize the amount of required storage.

The lung fish genome requires special processing due to its extreme size of the FASTA file of 86 Gigabytes (GB). This is described in the “Computational resources” section.

### Nullomer and sequence counting

With the genome data in a suitable format, the occurrences of nullomers or short sequences in general can be counted. The used algorithm is based on the description in the original paper about nullomers [1] with a few modifications. Here, we use a self-written code in the programming language python [40]. Array calculations were done with the library numpy [41], visualizations with library matplotlib [42] and just-in-time compilation as well as parallelization over multiple CPU cores was implemented using the library numba [43]. The latter is done to overcome the low computational speed of native python and achieves runtimes comparable to programming languages like C++ or Fortran which is necessary to perform the calculations with larger genomes at reasonable times.

Each sequence of a given sequence length can be represented in two different ways: Via an array containing entries corresponding to the base pairs as described in the section “Genome reading” or via a number. The latter representation can be calculated by treating the array representation as a number in base 4, leading to a single number which then can, for example, also be expressed in base 10. While the array representation is the direct output of genome files, the number representation has to be used to access arrays. Therefore, converting between these two representations is necessary in many situations and can be done straightforward via modular arithmetic.

The counting of nullomers is achieved by calculating the occurrences of all sequences of a given sequence length and then determining the number of sequences with zero occurrences. To compute the occurrences of each sequence, we iterate through the genome with a reading frame equal to the sequence length, incrementing by one base pair each step. The sequence in the reading frame can be transferred to a number. An array of size 4^*n*^ which documents the occurrence numbers of all sequences of length *n* is then incremented by one at the position of the computed number. To account for genomes with some undetermined base pairs, it is also checked that the sequence in the reading frame has no entry of –1 at any position.

This procedure can also be parallelized to run across multiple CPU cores. To do so, the genome is split into parts of the same length, with the number of parts being equal to the number of CPU cores. However, it has to be ensured that no race conditions occur if the array with the sequence counts is accessed by multiple threads at once. Therefore, this array should have a copy for each thread which are then summed up after all calculations have finished.

Next to counting sequences in the way which is described above, it is also necessary to consider the complementary DNA strand. This can be done without iterating through the genome again. Instead, the array with the stored sequence counts is considered. For each array index (corresponding to a sequence), the reverse complement of the sequence is computed which is transferred back to an index. At this index, the sequence count of the original index is added. If a lot of entries in the sequence counts array are zero (i.e. there are a lot of nullomers), it can also be checked if the array has a nonzero entry at the given index before computing the reverse complement sequence.

Note, that the above procedure is only faster than iterating through the genome for sequence lengths *n* where the total number of sequences (4^*n*^) is smaller than the genome length. Parallelization can be achieved by splitting up the genome. However, it is not necessary to create copies of the sequence counts array as there is no need to worry about race conditions, since every sequence has a unique reverse complement, meaning that each array index only gets accessed once.

After the sequence counts of both strands of the genome are computed in this way, the number of nullomers for each sequence length is equal to the amount of array entries which are zero. In addition to determining the nullomer count, the sequence count array is also stored for sequence lengths up to 13. The datatype “int32” is sufficient, except for the lung fish genome which needs “int64” for very short sequence lengths to avoid overflow.

For sequence lengths 14 and 15, the sequence counts are not stored due to their large sizes (about 1 and 4 GB, respectively). Also, at these sequence lengths, nullomers already make up a large fraction of the sequences, if not the majority for shorter genomes. If no sequence counts are stored, the parallelization scheme can be changed as well. Since the sequence count array sizes become quite large at lengths 14 and 15, creating multiple copies during the parallel calculations can lead to memory issues or slow down the calculations. Therefore, only one array is used, since the nullomer count—which is the only information that is stored at these sequence lengths—is not susceptible to race conditions.

This is the case, since the entries with nullomers are never accessed due to their sequence counts being zero. Consequently, no race conditions can occur in these entries. In addition to this, entries with counts larger than zero can also not become nullomers by accident: If an entry is accessed only once, no race condition can occur; if it is accessed more than once, a race condition can occur but only one increment is omitted, making the sequence count smaller than it should be but not zero. Furthermore, the occurrence of race conditions has been tested for the human genome, and only a few very common sequences rich in A and T exhibited lower sequence counts due to race conditions.

### Genome generation

Generating artificial genomes was done using a Markov chain model. To enable the genome generation, it is first necessary to compute the probabilities used in the Markov chain. This is done by iterating through the genome as described above for the sequence counting. However, this time the length of the reading frame is equal to the sum of the input and output lengths of a Markov chain (i.e. *n*+*m* for a Markov chain of the type *M*(*n, m*)). The sequence in the reading frame is then split into two smaller sequences with lengths *n* and *m* which are converted to indices separately. Beforehand, an array for the Markov chain probabilities is created with the dimensions 4^*n*^ and 4^*m*^. Now, the array is increased by one at the indices determined from the two sequences. After iterating through the genome, the sequence counts are normalized to convert them to probabilities.

Similarly, an array of probabilities has to be created for the first input in the Markov chain which does not have a preceding sequence. These probabilities can be computed as the normalized sequence counts with the sequence length equal to the input length of the Markov chain. Based on these precomputed probabilities, a Markov chain genome of a given length can be generated. The starting sequence is decided by selecting a random number between 0 and 1 and then choosing the sequence from the respective array based on that number. After that, at each step the input sequence of the Markov chain is read out from the part of the genome that is already generated and converted to a number. Another random number between 0 and 1 is chosen and the two-dimensional array of transition probabilities is read out based on the aforementioned computed sequence number and the random number. This results in a new output sequence which is added to the generated genome. The process is repeated until the generated genome has the given length.

### Computational resources

The genome generation and sequence counting for all species except for the lung fish can be done on a personal computer, due to the implementation with just-in-time compilation and parallelization. A bottleneck for the processing of reference genomes is the initial read from the FASTA files which need to be stored in array form. This takes longer than the sequence counting for the human genomes but cannot be avoided as an initial processing step.

In the case of the human and mouse genomes, the generation of ten genomes and the sequence counting took two hours on a personal computer, parallelized on 16 cores. The smaller genomes can be processed within minutes. For the lung fish genomes with a genome size of 86 GB (both for the FASTA sequence and if stored in “int8” format), the processing was done on a compute node and took around one day. The 232 human genomes were also processed on a compute node in around 1.5 days. During these calculations, the nullomer counts up to sequence lengths of 15 were stored as well as the sequence counts of all sequences up to length 13.

## Acknowledgements

Both authors acknowledge support from the Deutsche Forschungsgemeinschaft through the RTG 2749/1: “Biological Clocks on Multiple Time Scales”. Both authors would also like to acknowledge networking support by the COST Action CA21169, DYNALIFE, supported by COST (European Cooperation in Science and Technology). Calculations were performed on the Linux Cluster of the University of Kassel. L.W. and M.E.G. thank Katja Kapp for useful discussions.

## Author contributions

Luc Wieners: Conceptualization, Data curation, Formal analysis, Investigation, Methodology, Project administration, Software, Validation, Visualization, Writing – original draft, Writing – review & editing

Martin E. Garcia: Conceptualization, Funding acquisition, Investigation, Project administration, Resources, Supervision, Validation, Writing – review & editing

## Materials & Correspondence

Correspondence and requests for materials should be addressed to Luc Wieners.

